# *P. aeruginosa* rhamnolipids stabilize human rhinovirus 14 virions

**DOI:** 10.1101/2025.06.04.657910

**Authors:** Joshua J. Baty, Aidan K. Drozdick, Julie K. Pfeiffer

**Author notes:** Address correspondence to Julie K. Pfeiffer.

## Abstract

Many mammalian viruses encounter bacteria and bacterial molecules over the course of infection. Previous work has shown that the microbial ecology of the gut plays an integral role in poliovirus and coxsackievirus infection, where bacterial glycans can facilitate virus-receptor interactions, enhance viral replication, and stabilize viral particles. However, how airway bacteria alter respiratory viral infection is less understood. Therefore, we investigated whether a panel of airway bacteria affect rhinovirus stability. We found that *Pseudomonas aeruginosa*, an opportunistic airway pathogen, protects human rhinovirus 14 from acid or heat inactivation. Further investigation revealed that *P. aeruginosa* rhamnolipids, glycolipids with surfactant properties, are necessary and sufficient for stabilization of rhinovirus virions. Taken together, this work demonstrates that specific molecules produced by an opportunistic airway pathogen can influence a respiratory virus.

**Importance:** Bacteria can enhance viral stability and infection for enteric members of the *Picornaviridae* such as poliovirus and coxsackievirus; however, whether bacteria influence respiratory picornaviruses is unknown. In this study, we examined impacts of airway bacteria on rhinovirus, a major etiological agent of the common cold. We found that *P. aeruginosa* protects human rhinovirus 14 from both acid and heat inactivation through rhamnolipids. Overall, this work demonstrates bacterial effects on respiratory virus through specific bacterial molecules.

## Introduction

Rhinoviruses are the most common cause of the common cold (1-3). Rhinoviruses are a large and diverse group of enteroviruses that are divided into three species that bind various receptors—ICAM-1, LDLR, or CDHR3—that are found in the airway (2, 3). Although most rhinovirus infections are mild and self-limiting, severe and long-term consequences are possible. Rhinoviruses are the most common viral infection in those with cystic fibrosis and contribute to exacerbations (4-14). Cystic fibrosis disease is the result of ion imbalance at the cell surface (15), leading to aggregation of thick, sticky mucus and chronic colonization of opportunistic bacterial pathogens (16-18).

Previous work from our lab has shown that intestinal bacteria bind related enteroviruses such as poliovirus and coxsackievirus (19-23). Bacteria-virus interactions stabilize these viruses and protect from heat inactivation (19). Further, bacteria promote viral replication *in vivo*, as demonstrated by reduced titers of poliovirus and coxsackievirus in antibiotic-treated animals (22). Similarly, intestinal viruses in other families also benefit from bacteria, including mouse mammary tumor virus, murine norovirus, and certain strains of reovirus (21, 24-28). Although these interactions have been examined for these enteric viruses and related enteroviruses such as poliovirus and coxsackievirus, whether bacteria influence rhinovirus infections is unknown.

To determine if respiratory bacteria stabilize rhinovirus, we incubated human rhinovirus 14 (HRV14) with a panel of respiratory bacteria at an inactivating acidic pH of 5.8 or inactivating heat of 49°C and found that *P. aeruginosa*, a notorious cystic fibrosis pathogen, protects HRV14 from inactivation. Mechanistically, we found that rhamnolipids, biosurfactants produced by *P. aeruginosa*, are necessary and sufficient for this stabilization. Taken together, these results demonstrate that specific molecules from a ubiquitous bacterium can stabilize HRV14.

## Results

### *P. aeruginosa* stabilizes HRV14

Given that rhinoviruses likely encounter airway bacteria during infection, we questioned whether airway bacteria influence viral infection. In contrast to many other enteroviruses, rhinoviruses are acid sensitive (29). The healthy upper airway has an acidic pH that increases from the nares through the nose and sinuses with pHs of 5.5 to 6.5, respectively (30). The healthy lower airway has a neutral pH between 7.0-7.5 (31, 32). However, in the presence of inflammation, the respiratory tract pH can decrease.

During asthma exacerbations, exhaled breath condensate falls to 5.2 (33). In those with cystic fibrosis, exhaled breath condensate is reduced to 5.8 basally and to 5.3 during exacerbations (34). To examine potential effects of bacteria on HRV14 pH sensitivity, we first incubated 10^5^ PFU HRV14 in synthetic nasal media (35) at a pH of 6.8 or 5.8 for one hour before quantifying titer by plaque assay using H1 HeLa cells **(Fig 1AB)**. As expected, we found a >1000-fold reduction in viral titer at pH 5.8 (**Fig 1B**) compared with pH 6.8 (**Fig 1A)**. Next, we repeated the assay in the presence of a panel of airway bacteria. Many of the bacteria used in this screen (e.g. *M. catarrhalis, D. pigrum, S. aureus, S. epidermidis*) are commonly found in the upper airways from the nares to the sinuses (36, 37). However, some of these bacteria are enriched in the upper airways and colonize the lower airways during chronic pulmonary diseases such as cystic fibrosis (e.g. *P. aeruginosa, S. aureus, S. parasanguinis*)(38, 39). Overnight cultures of bacteria (10^6^-10^8^ CFU (**Table 1**)) were washed and resuspended in media at a pH of either 5.8 or 6.8, 10^5^ PFU HRV14 was added, and bacteria and virus were incubated together for one hour at 33°C prior to plaque assay. Bacteria had no effect on rhinovirus titers at a non-inhibitory pH of 6.8 (**Fig 1A**). At a pH of 5.8, HRV14 titers were reduced across all samples, with no bacterial strain significantly protecting HRV14 from acid inactivation (**Fig 1B**), although *P. aeruginosa* strains had increased yields that were not statistically significant in this initial broad screen. We repeated the pH 5.8 stability assay for HRV14 incubated with *P. aeruginosa* PAO1 and found that it significantly increased viral stability by 10-fold (**Fig 1B inset**).

**Table 1.** Bacterial strains.

| Strain | Characteristics | Overnight CFU/mL | Reference/Source |
| --- | --- | --- | --- |
| *P. aeruginosa* PAO1 | Wound, lab isolate | 10<sup>8</sup> | (57, 80) |
| PAO1 *rhIA* | Transposon insertion in *rhIA* | 10<sup>8</sup> | (57) |
| PAO1 *rhIB* | Transposon insertion in *rhIB* | 10<sup>8</sup> | (57) |
| PAO1 *rhIC* | Transposon insertion in *rhIC* | 10<sup>8</sup> | (57) |
| *P. aeruginosa* FRD1 | Cystic fibrosis, lab isolate | 10<sup>8</sup> | (81) |
| *P. aeruginosa* PA14 | Wildtype, lab isolate | 10<sup>8</sup> | (82) |
| *M. catarrhalis* | Wildtype | 10<sup>6</sup> | (83) |
| *E. coli* K12 | Lab isolate | 10<sup>8</sup> | (21) |
| *K. pneumoniae*<br>NCTC 9633 | Lab isolate | 10<sup>8</sup> | (84) |
| *D. pigrum* | Wildtype | 10<sup>6</sup> | (85) |
| *S. aureus* USA200 | MRSA | 10<sup>8</sup> | (86) |
| *S. aureus* Wichita | Wound, lab isolate | 10<sup>8</sup> | (87) |
| *S. epidermidis* | Wildtype | 10<sup>8</sup> | (88) |
| *S. parasanguinis*<br>FW213 | Wildtype | 10<sup>8</sup> | (89) |

**Figure 1.**
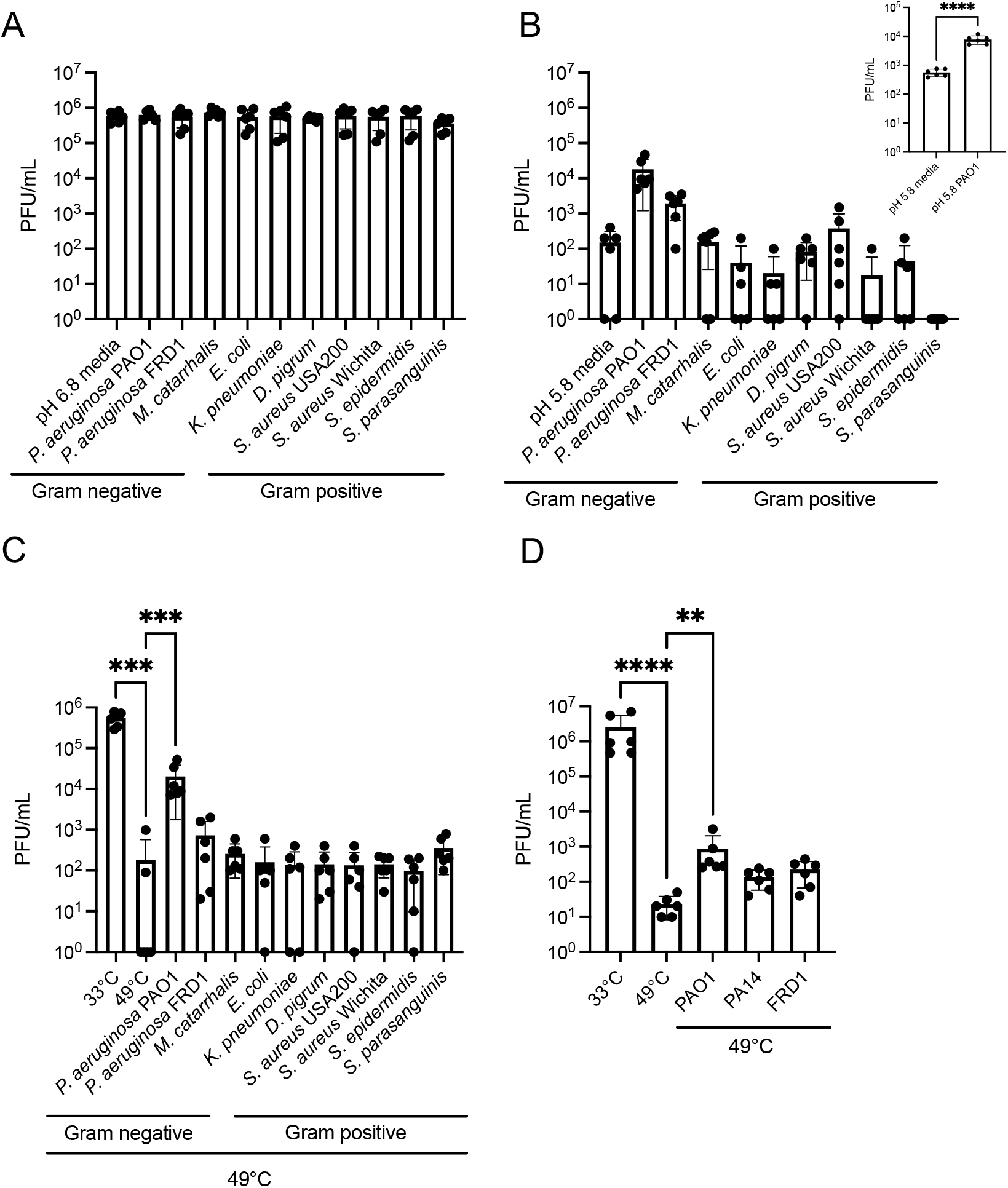
*P. aeruginosa* stabilizes HRV14. A/B/C) HRV14 (10^5^ PFU) was incubated for one hour at a pH of 6.8 (A) or 5.8 (B) or for two hours at 33°C or 49°C C) in the presence or absence of a panel of airway bacteria (10^6^-10^8^ CFU). Samples were centrifuged and PFU ere quantified from the supernatant by plaque assay. D) HRV14 was incubated in the presence or absence of. *aeruginosa* PAO1, PA14, or FRD1 at 33°C or 49°C for two hours prior to plaque assay. n=6, 3 biological eplicates with 2 technical replicates. **, p<0.01, ***, p<0.001, ****, p<0.0001 (A-D Kruskal-Wallis, Dunnett’s ost hoc test, B insert unpaired t test).

We next determined whether airway bacteria could protect HRV14 from heat inactivation. For these experiments, HRV14 was incubated at 33°C or 49°C for two hours, followed by titer analysis via plaque assay. As expected, HRV14 titers were reduced by >1000-fold after incubation at 49°C (**Fig 1C**). In the presence of airway bacterial strains, only *P. aeruginosa* PAO1 significantly increased HRV14 titers at 49°C (**Fig 1C**). We next examined whether increased HRV14 viability in the presence of *P. aeruginosa* was unique to the strain PAO1 or if other strains of *P. aeruginosa* conferred protection. We compared *P. aeruginosa* strains PAO1, PA14, and FRD1. All three of these strains are typical lab strains of *P. aeruginosa*; however, exopolysaccharide and virulence factor production vary (40-42). We found that the PAO1 strain significantly increased HRV14 recovery after heat exposure, but PA14 and FRD1 strains did not (**Fig 1D**), suggesting that PAO1 stabilizes HRV14 more than other strains of *P. aeruginosa*.

### HRV14 does not have increased binding to *P. aeruginosa*

Our group previously reported that direct binding to bacteria and bacterial glycans stabilizes related picornaviruses such as poliovirus and coxsackievirus (19-22, 43). Therefore, to determine whether *P. aeruginosa* PAO1 has increased binding to HRV14, potentially explaining its virion stabilization phenotype, we quantified binding of purified, ^35^S-radiolabeled HRV14 to bacterial strains. We incubated our panel of airway bacteria (**Table 1**) with ^35^S-radiolabeled HRV14 (4,000 CPM/10^6^ PFU) at pHs of 5.8 or 6.8 to determine if HRV14 binds relevant airway bacteria. Virus was also incubated with 2.8 µm streptavidin beads to account for nonspecific binding. *E. coli* and *S. aureus* Wichita had significantly increased HRV14 binding compared to the bead control at a pH of 6.8 (**Fig 2A**), although no significant differences in binding were observed at Ph 5.8 (**Fig 2B**). Surprisingly, HRV14 did not display enhanced binding to *P. aeruginosa*, suggesting that direct binding may not be a major facet of stabilization against acid inactivation.

**Figure 2.**
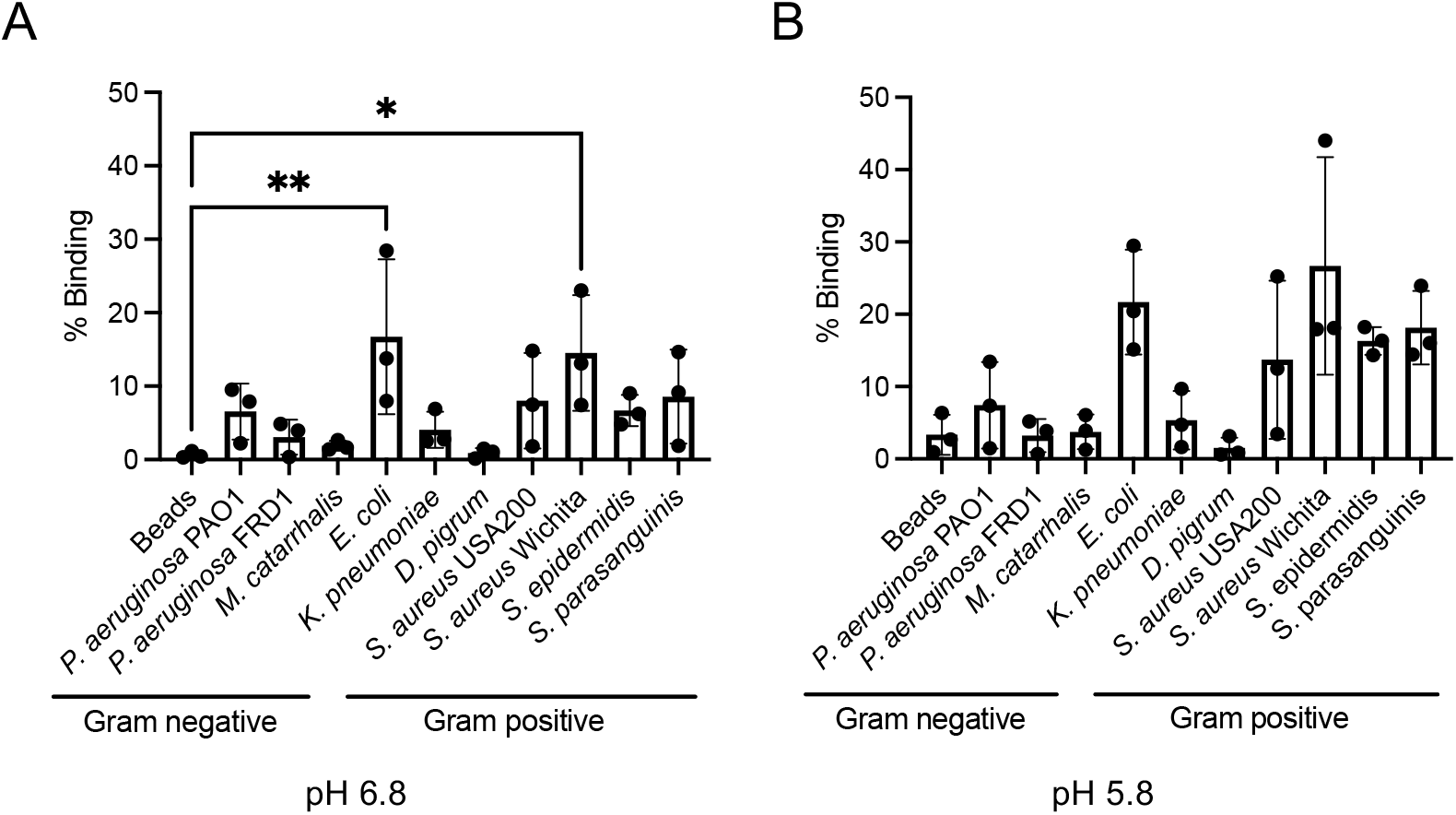
HRV14 does not have enhanced binding to *P. aeruginosa*. A/B) ^35^S-radiolabeled HRV14 (∼4,000 CPM/ 10^6^ PFU) was incubated in the presence or absence of streptavidin ds (2.8 µm) or 10^6^-10^8^ CFU bacteria in media at a pH of 6.8 (A) or 5.8 (B) at 33°C for one hour. Samples were trifuged and washed to remove unbound virus. Bound virus was quantified via scintillation counting and malized to input. n=3. A) ns, p>0.05 (Kruskal-Wallis, Dunnett’s post hoc test). B) *, p<0.05, **, p<0.01 (one-way OVA, Dunnett’s post hoc test).

### Heat-killed *P. aeruginosa* stabilizes HRV14

To determine if *P. aeruginosa*-mediated protection of HRV14 from acid and heat inactivation was due a heat-sensitive factor or relied upon active *P. aeruginosa* metabolism, HRV14 was incubated with live or heat-killed *P. aeruginosa* at a pH of 6.8 vs. 5.8 (**Fig 3A**) or at 33°C vs. 49°C (**Fig 3B**). Heat-killed *P. aeruginosa* protected HRV14 from acid inactivation, suggesting that a heat stable *P. aeruginosa* factor stabilizes HRV14 (**Fig 3A**).

**Figure 3.**
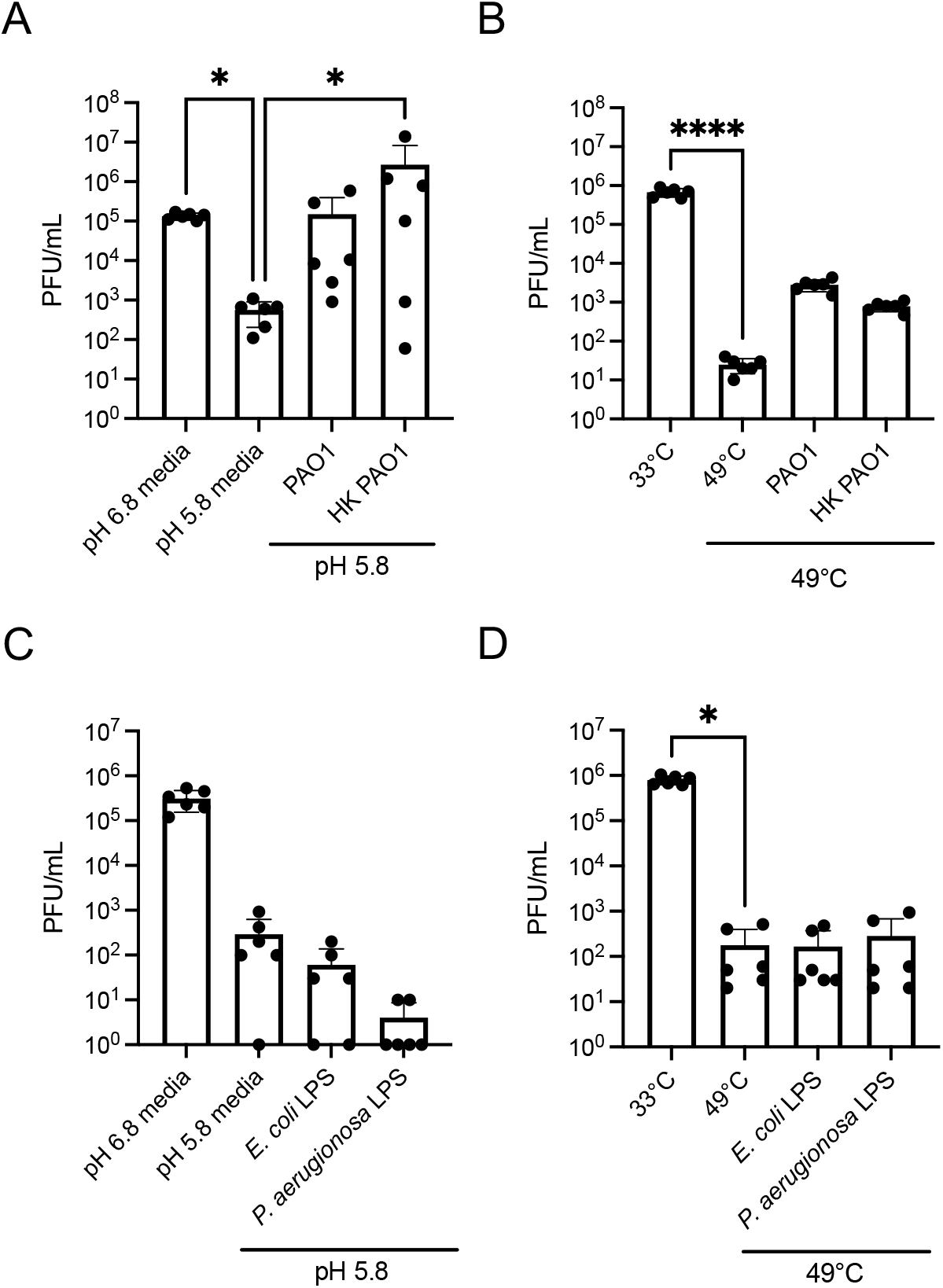
Heat-killed *P. aeruginosa* stabilizes HRV14. A/B) HRV14 (10^5^ PFU) was incubated in the presence or absence of 10^8^ CFU live or heat-killed (HK) *P. eruginosa* PAO1 at a pH of 5.8 or 6.8 at 33°C for one hour (A) or 33°C or 49°C for two hours (B) prior to laque assay. C/D) HRV14 was incubated in the presence or absence of 1 mg/ml LPS from *E. coli* or *P. eruginosa* at a pH of 5.8 or 6.8 for one hour (C) or 33°C or 49°C for two hours (D) prior to plaque assay. n=6, biological replicates with 2 technical replicates. *, p<0.05, ****, p<0.0001 (A/C/D, Kruskal-Wallis, Dunnett’s ost hoc test, B, one way ANOVA, Dunnett’s post hoc test).

Given that heat-killed *P. aeruginosa* was sufficient to protect HRV14 from acid inactivation, and our past work demonstrated that heat stable bacterial lipopolysaccharide (LPS) stabilizes picornaviruses, we hypothesized that LPS stabilizes HRV14. As an external glycan moiety on Gram-negative bacterial surfaces, LPS is a common factor that rhinovirus is likely to encounter. Previous work from our lab demonstrated that poliovirus binds LPS and that binding to LPS stabilizes poliovirus, Aichivirus, and coxsackievirus (19, 22). Conversely, LPS destabilizes enveloped influenza virions as well as alphavirus and flavivirus virions (44, 45). To assess LPS effects, HRV14 was incubated with LPS isolated from *E. coli* or *P. aeruginosa* at a pH of 5.8 vs. 6.8 for one hour (**Fig 3C**) or at 33°C vs. 49°C for two hours (**Fig 3D**) followed by plaque assay. Surprisingly, LPS did not protect HRV14 from acid or heat, suggesting that some other *P. aeruginosa* factor is responsible for stabilization.

### Rhamnolipids stabilize HRV14

We next hypothesized that other heat-stable, high abundance *P. aeruginosa* molecules stabilize HRV14. Like LPS, rhamnolipids are glycolipids that are produced by *P. aeruginosa* at high concentrations (46, 47). Rhamnolipids are important for biofilm formation and architecture, motility, and protection from phagocytosis (48-51). Rhamnolipids are synthesized by the enzymes RhlA, RhlB, and RhlC (**Fig 4A**)(52). RhlA catalyzes the conversion of B-hydroxyacyl-ACP into the fatty acid dimer 3-(3-hydroxyalkanoyloxy)alkanoates (HAA)(53, 54). RhlB is a rhamnosyltransferase that catalyzes a reaction between HAA and dTDP-L-rhamnose to produce mono-rhamnolipids (55). RhlC acts as a second rhamnosyltransferase that catalyzes the conversion of mono-rhamnolipids and dTDP-L-rhamnose to di-rhamnolipids (56).

**Figure 4.**
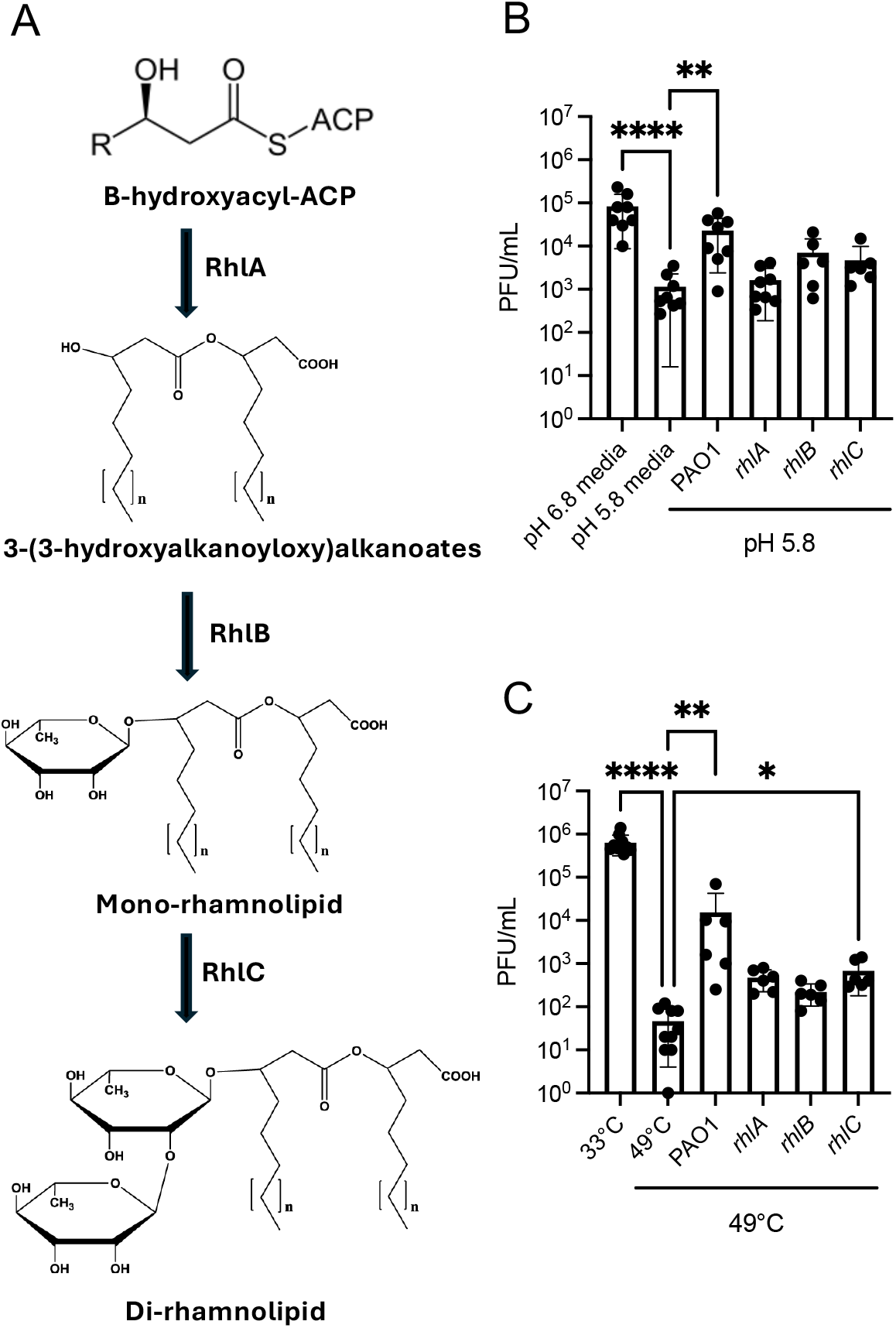
Insertion mutation of rhamnolipid synthesis genes ablates HRV14 stabilization. A) *P. aeruginosa* rhamnolipid synthesis pathway. B/C) HRV14 (10^5^ PFU) was incubated in the presence or bsence of 10^8^ CFU PAO1, *rhlA, rhlB*, or *rhlC* transposon insertion mutants at a pH of 5.8 or 6.8 at 33°C for one our (B) or 33°C or 49°C for two hours (C) prior to plaque assay. n=6-8, 3-4 biological replicates with 2 technical eplicates. *, p<0.05, ** p<0.01, ****, p<0.0001 (Kruskal-Wallis, Dunnett’s post hoc test).

To examine the potential impact of *P. aeruginosa* rhamnolipids on HRV14 stabilization, we used strains with transposon insertions within rhamnolipid synthesis genes to test for necessity, and addition of purified rhamnolipids to test for sufficiency. First, we obtained *rhlA, rhlB*, and *rhlC* mutants in the *P. aeruginosa* PAO1 background (57) and incubated them with HRV14 at a pH of 5.8 vs. 6.8 for one hour (**Fig 4B**) or at 33°C vs. 49°C for two hours **(Fig 4C)** followed by plaque assay. All mutants in the rhamnolipid synthesis pathway failed to protect HRV14 from acid inactivation (**Fig 4B**). The *rhlC* mutant partially restored protection of HRV14 from heat inactivation, suggesting that the production of mono-rhamnolipids are somewhat protective against heat (**Fig 4C**). Next, we tested whether purified rhamnolipids could stabilize HRV14 in the absence of bacteria. HRV14 was incubated with various concentrations of rhamnolipids at a pH of 5.8 vs. 6.8 for one hour (**Fig 5A**) or at 33°C vs. 49°C for two hours **(Fig 5B)** followed by plaque assay. Rhamnolipids protected HRV14 from both acid and heat inactivation at a concentration of 0.5 mg/mL. Overall, data in Figures 4 and 5 indicate that rhamnolipids are necessary and sufficient for stabilization of HRV14 by *P. aeruginosa*.

**Figure 5.**
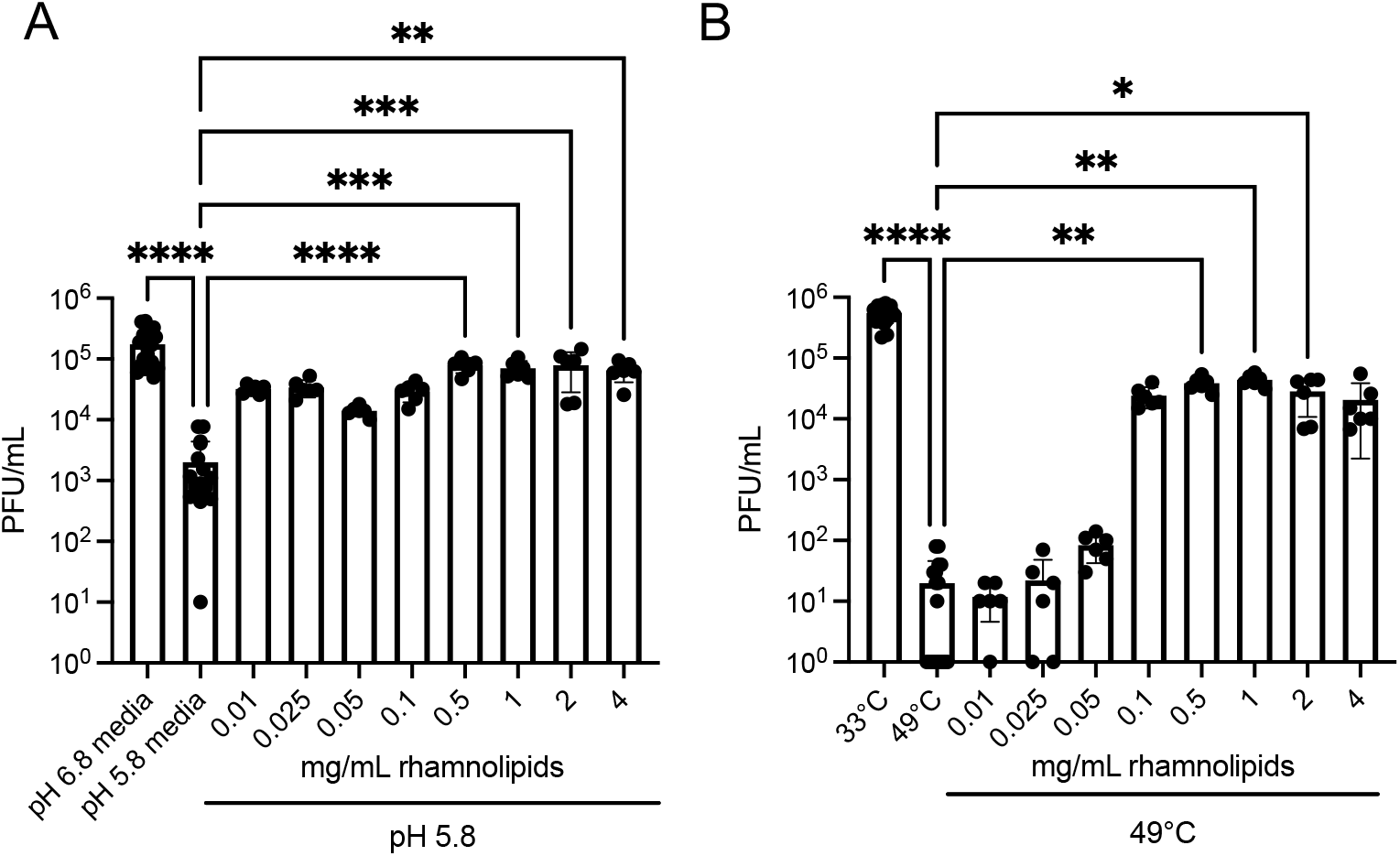
Rhamnolipids stabilize HRV14. A/B) HRV14 (10^5^ PFU) was incubated in the presence or absence of various concentrations of rhamnolipids at a of 5.8 or 6.8 at 33°C for one hour (A) or 33°C or 49°C for two hours (B) prior to plaque assay. n=6, 3 biological plicates with 2 technical replicates. *, p<0.05, ** p<0.01, ***, p<0.001 ****, p<0.0001(Kruskal-Wallis, Dunnett’s st hoc test).

To confirm that rhamnolipids stabilize HRV14 using an assay independent from viral viability assays, we performed a cell-free <u>Pa</u>rticle <u>S</u>tability <u>T</u>hermal <u>R</u>elease assa<u>y</u> (PaSTRy)(58). Through this assay, virion RNA release is measured over a temperature gradient using SYBR green II dye to define the exact temperature of virion inactivation. RNA release was measured for HRV14 in the presence or absence of *P. aeruginosa* LPS (as a negative control) or rhamnolipids (**Fig 6**). Untreated HRV14 released RNA at 48.7°C. As expected from our plaque-based assays, LPS had no effect on the temperature at which HRV14 RNA release occurred. However, rhamnolipids shifted HRV14 RNA release temperatures by ∼1°C at 0.05 and 0.1mg/mL concentrations and by ∼3°C at 1mg/mL concentration. Taken together, these results demonstrate that rhamnolipids stabilize HRV14.

**Figure 6.**
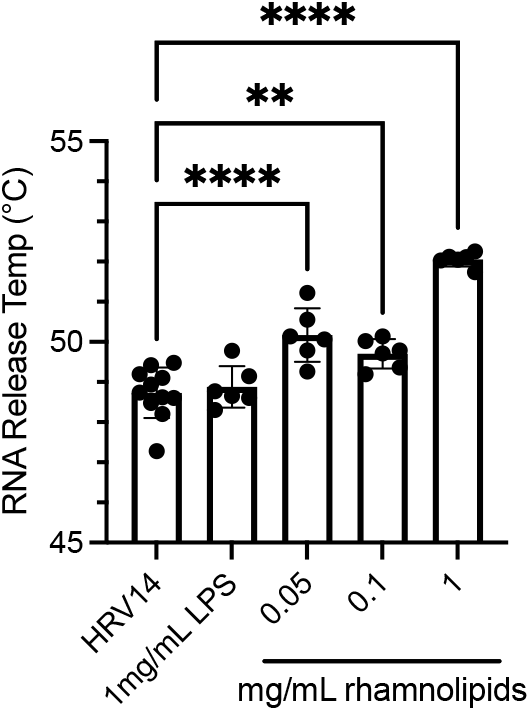
Rhamnolipids enhance HRV14 thermostability. HRV14 thermostability profile using a cell-free Particle Stability Thermal Release assay (PaSTRy). HRV14 (10^5^ PFU) s added to SYBR green II with or without LPS or rhamnolipids. Samples were heated from 25°C to 95°C on a 1% pwise gradient with fluorescence monitoring. n=6, 3 biological replicates with 2 technical replicates. **, p<0.01, *, p<0.0001 (one-way ANOVA, Dunnett’s post hoc test).

## Discussion

Rhinoviruses are important respiratory pathogens, but potential impacts of bacteria on rhinovirus infection are largely unknown. Here, we screened a panel of respiratory bacteria and found that *P. aeruginosa*, an opportunistic pathogen that establishes chronic infections in those with chronic airway diseases, protects HRV14 from acid and heat inactivation. Investigation of *P. aeruginosa* strains deficient for rhamnolipid production and addition of exogenous rhamnolipids revealed that rhamnolipids were necessary and sufficient for HRV14 stabilization.

Rhamnolipids are glycolipids that are important for *P. aeruginosa* physiology and infection. *P. aeruginosa* produces copious amounts of rhamnolipids, with wildtype *P. aeruginosa* PAO1 producing as much as 39 g/L (46). Critically, rhamnolipids are present in sputum samples from people with cystic fibrosis that are colonized with *P. aeruginosa* (59, 60). Rhamnolipids help shape biofilm architecture (49), mediate *P. aeruginosa* dispersal (61-63), enhance *P. aeruginosa* motility (62), decrease phagocytosis (50, 51), and damage cell membranes (51, 64). Rhamnolipids inhibit the colonization and disperse a wide array of other bacteria (63, 65-70). Additionally, rhamnolipids inactivate enveloped viruses such as herpesviruses, coronaviruses, and respiratory syncytial virus via envelope disruption (71-75).

Less is known about interactions between rhamnolipids and nonenveloped viruses, such as rhinoviruses and other picornaviruses. Rhamnolipids have no effect on poliovirus stability (71), but *in silico* modeling of HRV14 suggested that rhamnolipids interact with the canyon region of the capsid, where rhinoviruses bind their receptors (76). This interaction may be responsible for the stabilization phenotype herein, but further studies are required to fully delineate the role of rhamnolipids during rhinovirus infection. Beyond viral stability, exposure to biosurfactants, such as rhamnolipids, can increase pathogenesis of other picornaviruses, such as encephalomyocarditis virus (EMCV)(77-79). The pesticides dichloro-diphenyl-trichloroethane (DDT) and fenitrothion are associated with clusters of Reye’s Syndrome, a rare condition involving liver pathology and brain swelling that often accompanies viral infection. These surfactants increase EMCV uncoating in treated cells (78). Additionally, these compounds reduce interferon responses, contributing to increased morbidity and mortality in mice (78).

Taken together, we found that rhamnolipids, glycolipids produced by the opportunistic pathogen *P. aeruginosa*, increases stability of HRV14. This interaction may be clinically relevant as many people with cystic fibrosis are chronically colonized by *P. aeruginosa* and rhinoviruses are a common cause of exacerbation events. Future studies are necessary to determine the role rhamnolipids and other compounds play over the course of rhinovirus infection.

## Methods

### Cells and viruses

HeLa H1 cells were propagated in DMEM supplemented with 10% calf bovine serum and 1% antibiotics. Cells were grown at 37°C with 5% CO_2_. HRV14 was propagated from an infectious clone (gift of William Jackson) and infections were performed at 33°C 5% CO_2_.

### Bacterial strains, culture conditions, and reagents

*P. aeruginosa* PAO1, FRD1, PA14 and the PAO1 isogenic mutants, *rhlA, rhlB, rhlC, S. aureus, S. epidermidis, E. coli*, and *K. pneumoniae* were maintained on lysogeny broth (LB) agar and grown on LB at 37°C with shaking at 250 rpm. *M. catarrhalis* was grown in brain heart infusion media at 37°C with shaking at 250 rpm. *D. pigrum* and *S. parasanguinis* were grown in Todd Hewitt broth/agar at 37°C with 5% CO_2_. Synthetic nasal media was prepared as described in Krismer et al 2014 (35).

### Quantifying effects of bacteria on viral stability

For acid sensitivity assays, overnight cultures of bacteria (**Table 1**, 10^6^ or 10^8^ CFU, depending on the strain) were centrifuged, washed in synthetic nasal media (SNM) at a pH of either 5.8 or 6.8, centrifuged and resuspended in (SNM) at a pH of either 5.8 or 6.8. HRV14 (10^5^ PFU) was added, and the virus and bacteria of interest were incubated at 33°C with 5% CO_2_ for one hour. For heat sensitivity assays, overnight cultures were centrifuged, washed with PBS, centrifuged and resuspended in PBS. HRV14 was added as above, and the mixture was incubated at either 33°C or 49°C for two hours. After each incubation, samples were centrifuged and PFU in the supernatants were quantified via plaque assay as described (19). Briefly, samples were diluted in PBS supplemented with 100 µg/mL CaCl_2_ and 100 µg/mL MgCl_2_ and allowed to attach to cells for 30 minutes at 33ºC with 5% CO_2_. Agar overlays containing DMEM with 10% calf bovine serum and 1% antibiotics was added and removed 48 hours after infection. PFU were enumerated following crystal violet staining of monolayers.

### Quantifying viral binding to bacterial cells

^35^S-radiolabeled HRV14 was generated as previously described (20). Briefly, infected cells were pulsed with ^35^S-amino acids to label progeny virions, cell-associated virions were collected, and purified using Capto Core 700 beads (Cytivia) according to the manufacturer’s instructions. Briefly, rhinovirus was mixed end-over-end at 4°C with capto core beads for 45-minute increments three times. The slurry was centrifuged and virus from the supernatant was assessed for purity by SDS-PAGE. For binding assays, ∼4000 counts per minute (CPM)(10^6^ PFU) HRV14 was added to overnight bacterial cultures or streptavidin beads (Invitrogen, Dynabeads) resuspended in SNM pH 5.8 or 6.8. Incubation proceeded for one hour at 33ºC with 5% CO_2_ and the mixture was centrifuged and washed to remove unbound virus. The pellet was resuspended in Budget-Solve complete counting cocktail (Research Products International) and CPM was determined by scintillation counting.

### Quantifying effects of lipopolysaccharide and rhamnolipids on viral stability

Live or heat-killed *P. aeruginosa* PAO1 was incubated with HRV14 as above.

PAO1 was heat-killed by incubating at 95ºC for 10 minutes. LPS (at 1 mg/mL) from *E. coli* (O111:B4, Sigma) or *P. aeruginosa* (PA-10, Sigma) was resuspended in SNM Ph 5.8 or PBS and incubated at 33ºC or 49ºC and quantified via plaque assay as above.

Exogenous rhamnolipids from *P. aeruginosa* (Sigma) were added and incubated via the same scheme.

### Particle Stability Thermal Release assay (PaSTRy)

Capto-core (Cytiva) purified and Amicon filter-concentrated (Sigma) HRV14 (∼10^5^ PFU) was combined with rhamnolipids, SYBR green II (10x final concentration, Invitrogen), and buffer (10mM HEPES at pH 8, 200mM NaCl). The 50 uL reactions were heated from 25ºC to 95ºC on a 1% gradient in an ABI 7500 real-time thermocycler (Applied Biosystems) with fluorescent monitoring.

## Data analysis

All statistical analyses were performed using GraphPad Prism version 10.4.2 for macOS. Normality was assessed via the Shapiro-Wilk test. Further analyses were performed where indicated.

## Acknowledgements

This work was funded through NIH R37AI074668 to JKP. JJB was funded through NIH T32 AI007520. The *P. aeruginosa* rhamnolipid mutants were obtained from University of Washington Pa two-allele transposon mutant library, supported by The Cystic Fibrosis Foundation, grant SINGH24R0. We thank Dr. Tamia Harris-Tryon for the *Staphylococcus* isolates and Dr. Arielle Woznica for rhamnolipids.

